# Nano Random Forests to mine protein complexes and their relationships in quantitative proteomics data

**DOI:** 10.1101/050302

**Authors:** Luis F. Montaño-Gutierrez, Shinya Ohta, Georg Kustatscher, William C. Earnshaw, Juri Rappsilber

## Abstract

The large and ever-increasing numbers of quantitative proteomics datasets constitute a currently underexploited resource for drawing biological insights on proteins and their functions. Multiple observations by different laboratories indicate that protein complexes often follow consistent trends. However, proteomic data is often noisy and incomplete–members of a complex may correlate only in a fraction of all experiments, or may not be always observed. Inclusion of potentially uninformative data hence imposes the risk of weakening such biological signals. We have previously used the Random Forest (RF) machine-learning algorithm to distinguish functional chromosomal proteins from ‘hitchhikers’ in an analysis of mitotic chromosomes. Even though it is assumed that RFs need large training sets, in this technical note we show that RFs also are able to detect small high-covariance groups, like protein complexes, and relationships between them. We use artificial datasets to demonstrate the robustness of RFs to identify small groups even when working with mixes of noisy and apparently uninformative experiments. We then use our procedure to retrieve a number of chromosomal complexes from real quantitative proteomics results, which compare wild-type and multiple different knock-out mitotic chromosomes. The procedure also revealed other proteins that covary strongly with these complexes suggesting novel functional links. Integrating the RF analysis for several complexes revealed the known interdependency of kinetochore subcomplexes, as well as an unexpected dependency between the Constitutive-Centromere-Associated Network (CCAN) and the condensin (SMC 2/4) complex. Serving as negative control, ribosomal proteins remained independent of kinetochore complexes. Together, these results show that this complex-oriented RF (nanoRF) can uncover subtle protein relationships and higher-order dependencies in integrated proteomics data.

**Abbreviations**
RF: Random Forest
MCCP: Multi-Classifier Combinatorial Proteomics
nanoRF: Random forests trained with small training sets
MVP: Multivariate proteomic profiling
FP: Fractionation profiling
ICP: interphase chromatin probability
CCAN: Constitutive Centromere-Associated Network
Nup: Nucleoporin
SMC: Structural Maintainance of Chromosomes
SILAC: Stable Isotope Labeling by Amino acids in Cell culture

## INTRODUCTION

Proteins influence many processes in cells, often affecting the synthesis, degradation and physicochemical state of other proteins. One strategy that diversifies and strengthens protein functions is the formation of multi-protein complexes. For this reason, identification of partners in complexes is a powerful first step to determining protein function. However, determination of membership to or interaction with protein complexes remains an arduous task, mainly achieved via demanding biochemical experimentation. The latter can be limited by the ability to overexpress, purify, tag, stabilize, and obtain specific antibodies for the proteins in complexes of interest. Thus, any methods that facilitate protein complex identification and monitoring (1-3) have the potential to accelerate the understanding of biological functions and phenotype. The vast amount of proteomics data already available represents a largely untapped resource that could potentially reveal features currently undisclosed by traditional analysis, such as condition-dependent links, inter-complex contacts and transient interactions.

To date, co-fractionation is the gold standard to prove membership of protein complexes. This is based on the fact that proteins with the same mass, charge, elution rate, etc. will be part of the same fraction-i.e. co-fractionate– in techniques such as chromatography or gel electrophoresis. Yet, even in ideal cases, spurious proteins will co-fractionate with (contaminate) the complex of interest(4). One way to distinguish bona-fide members is to combine several fractionation experiments, as well as perturbations(5). Members of a complex will behave coordinately, whereas contaminants will usually behave more randomly. From a quantitative perspective, this translates into protein covariance - the covariance of proteins within a complex is stronger than that among contaminants. As additional biochemical fractionation conditions are considered, high covariance sets true members of a complex apart from contaminants or hitchhikers. This principle has been used recently in a large-scale effort that predicted 622 putative protein complexes in human cells by assessing the coordinated behaviour of proteins across several fractionation methods, among others (Havugimana et al., 2012; Michaud et al., 2012;).

Covariance among members of protein complexes has been observed in several integrative proteomics experiments (8, 9) and even used to predict association with complexes (8, 10). This relies on the fact that the co-fractionation of proteins that are functionally interconnected will be affected by common parameters, such as knock-outs or varying biochemical purification conditions. However, performing covariance analysis using multiple quantitative proteomics datasets is non-trivial. First, experimental or biological noise hampers quantitation of protein levels. Second, only a fraction of the experiments may be informative for any given complex. Third, proteins may go undetected, leading to missing values. Fourth, the relationship between different protein groups may only be observed under specific circumstances. The power of multivariate analysis methods like Principal Component Analysis (PCA), hierarchical clustering or k-nearest neighbours could be limited when a protein complex’s signal in the data is affected in all these ways. Here we show that the supervised machine learning technique Random Forests can overcome these limitations, distinguish the covariance of small protein groups, and provide biologically sound, predictive insights to protein complex composition, relationships and function. We describe this approach using as an example the behaviour of multi-protein complexes in mitotic chromosomes.

## EXPERIMENTAL PROCEDURES

### Cell Culture

As reported in (11), DT40 cells with wild-type genes (clone 18), as well as conditional knockouts for SMC2, CAP-H, CAP-D3, Scc1, or SMC5 were maintained in Roswell Park Memorial Institute (RPMI) 1640 medium (Wako Pure Chemical Industries Ltd.) supplemented with 10% (v/v) fetal bovine serum (FBS), 1% calf serum, 100 U/mL penicillin, and 100 μg/mL streptomycin (Wako Pure Chemical Industries Ltd.) at 39°C in a humidified incubator with an atmosphere containing 5% CO_2_ (12-15); For ^13^C and ^15^N labeling of lysine and arginine, cells were maintained in RPMI without L-lysine and L-arginine (Thermo Fisher Scientific, Waltham, MA, USA) supplemented with 10% (v/v) FBS dialyzed against a 10,000-molecular-weight cut-off membrane (Sigma-Aldrich, St. Louis, MO, USA), 100 μg/mL ^13^C_6_, ^15^N_2_-L-lysine: 2HCl, 30 μg/mL ^13^C_6_, ^15^N_4_-L-arginine: HCl (Wako Pure Chemical Industries Ltd.), 100 U/mL penicillin, and 100 μg/mL streptomycin (Gibco-BRL; Thermo Fisher Scientific) at 37°C in a humidified incubator with an atmosphere containing 5% CO_2_. To generate SMC2^OFF^, CAP-H^OFF^, CAP-D3^OFF^, Scc1^OFF^, or SMC5^OFF^ cells, SMC2^ON/OFF^, CAP-H^ON/OFF^, CAP-D3^ON/OFF^, Scc1 ^OFF^, or SMC5 ^OFF^ cells were grown in the presence of doxycycline for 30, 26, 24, 19, or 60 h, respectively, prior to blocking with nocodazole to inhibit expression. HeLa and U2OS cells in the exponential growth phase were seeded onto coverslips and grown overnight in Dulbecco‘s Modified Eagle’s Medium (DMEM) supplemented with 10% FBS at 37°C in an atmosphere containing 5% CO_2_.

### Mitotic chromosome isolation and SILAC

DT40 cells were incubated with nocodazole for 13 h, resulting in a mitotic index of 70%-90%. Mitotic chromosomes were isolated using a polyamine-ethylenediaminetetraacetic acid buffer system optimized for chicken DT40 cells (16). Five OD_260_ units were obtained from pooling the material of 4 independent preparations totaling 1.0 × 10^9^ DT40 cells and solubilized in sodium dodecyl sulfate-polyacrylamide gel electrophoresis (SDS-PAGE) sample buffer. Mitotic chromosomes of wild type and knockout cell lines were mixed in equal amounts judging by Picogreen quantification, except for the Ska3 KO experiment (9)where samples were equated using Histone H4 as a reference.

### Mass-spectrometric analysis

Proteins were separated into high-and low-molecular weight fractions by SDS-PAGE, in-gel digested using trypsin (17), and fractionated into 30 fractions each using strong cation-exchange chromatography (SCX). The individual SCX fractions were desalted using StageTips (18)and analyzed by liquid chromatography-MS on a LTQ-Orbitrap (Thermo Fisher Scientific) coupled to high-performance liquid chromatography via a nanoelectrospray ion source. The 6 most intense ions of a full MS acquired in the Orbitrap analyzer were fragmented and analyzed in the linear-ion trap. The MS data were analyzed using MaxQuant 1.0.5.12 for generating peak lists, searching peptides, protein identification (19), and protein quantification against the UniProt database (release 2013_07).

### Preparation of MS data for nanoRF

The SILAC ratios from the ‘Protein groups’ Maxquant output table were used directly. As for the Ska3 knock out experiment, SILAC ratio column values were directly taken from (9), and re-indexed according to the rest of the experiments. The ratio columns in table S1 were directly used for the analysis. All the raw MS and Maxquant output data, including those from the Ska3 experiment (9) via ProteomeXchange with identifier PXD003588. Missing values were substituted by the median value of each experiment, as is common practice in Random Forest applications. We reasoned that doing so would penalize the lack of observations by giving the same score to missing proteins of both positive and negative classes, which in turn increases the intersection between classes and thereby impacts separation quality.

### Random Forest analysis

The analysis was done with a custom R pipeline based on the Random Forests algorithm of Leo Breiman and Adele Cutler's Random Forest^™^ algorithm (20), implemented in R (21). All our scripts used are freely available through a Github repository (22) and include a step-by-step R guide script to perform nanoRF on any particular dataset. The RF algorithm attempts to find a series of requirements in the data that are satisfied by the positive training class and not by the negative training class. All these decisions are performed sequentially, hence they become a decision tree. An example of a decision tree would be “proteins with values >x in experiments 1 and 2. Out of those, proteins with values < y in experiments 3 and 5”. As the best set and decision sequence is not known a priori, the best bet is to generate many decision trees at random (hence the name random forest). Each tree votes for all compliant proteins as members of the positive class. The clearer the difference between the two classes in the data, the larger the number of trees that will vote for the positive class as indeed positive. The RF score (calculated for each protein) is the fraction of trees that voted for a protein as positive. In order to get a score for the members of the positive class as well, during the generation of each tree, some of the members of the positive and negative class are left out and treated as unknown. This Out-of-bag (OOB) procedure intrinsically controls for training set bias.

We set the number of trees in the forest to 3000 in each run. The Matthews correlation coefficient was calculated by using the formula

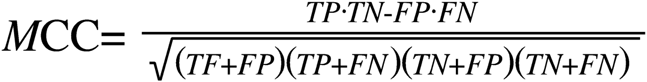

where TP indicates true positives, FP false positives, TN true negatives and FN false negatives. For null values of any of the sums in the denominator, the MCC was defined as 0. To choose a particular RF-Score as a cut-off, we evaluated 100 possible cut-offs between RF-scores 0 and 1 and kept that which maximized the MCC. In for cutoffs with the same maximum MCC, the smallest RF was chosen as a cut off to maximize sensitivity. Table S2 was directly used for machine learning.

### Informative experiment fraction VS noise analysis

We arbitrarily generated 600 matrices with ~5000 ‘protein’ rows and 20 ‘experiment’ columns (sizes similar to our SILAC ratio matrix) by sampling a standard normal distribution. In each matrix, 365 ‘proteins’ were selected to be part of the negative set and 5 groups of 12 proteins were set to be identical within their group in 2 … 20 ‘experiments’ (Figure 1D-F, horizontal axis). Next, Gaussian noise with standard deviation of .02 … 2 was added to the entire matrix (Figure 1D-F, vertical axis). Missing values were not added to the simulations as the RF pipeline would only transform NAs into the median value of the experiment and therefore just have the same effect as noise addition. RF analysis was then run for the 5 groups versus the negative set. Lastly, we calculated the mean of means of the RF scores for each positive group. The correlation was the mean of intra-group correlations of all positive groups.

**Figure 1.**
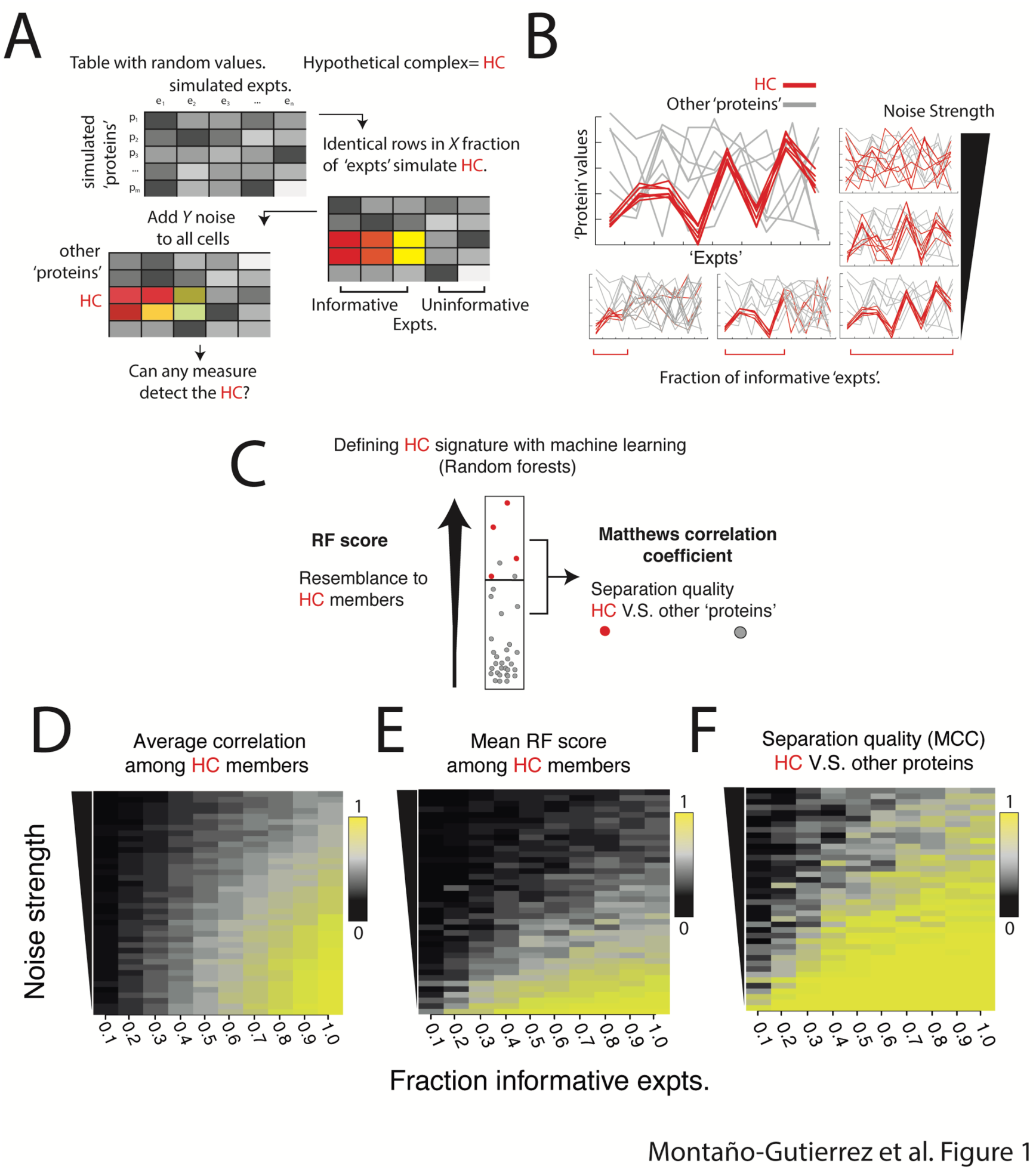
Supervised Machine Learning algorithm Random Forests can detect small, correlated protein groups in artificial proteomics data. A. Depiction of the procedure used to simulate proteomics data with ‘protein’ rows and ‘experiment’ columns. Some rows are made identical (red tones) in a fraction of experiments to simulate a hypothetical complex (HC), and Gaussian noise is then added element-wise to each table entry. B. Visual description of a hypothetical complex (red) versus other randomly generated proteins (grey) as the number of experiments (left-right) and the noise (bottom-up) affect the protein values in the experiments (all subpanels). C. Diagram to visualize the output from machine learning technique Random Forests. The RF score denotes the resemblance to the complex, while separation quality indicates how easily unrelated proteins covary with the complex. Red and grey dots depict the hypothetical complex and other proteins respectively. D,E,F. heat maps showing how the fraction of informative experiments (X axis) and the noise amount (Y axis) affect the Mean correlation (D) Random Forest score (E) and separation quality (F) of proteins in a complex. In each square, the value projected is the mean of means of 5 independent groups.

### Definition of protein group covariance

The covariance between random variables is only defined pairwise, and as such, the ‘mean correlation of a complex’ as mentioned in the text could be seen as a matrix A where A_ij_ is the correlation of protein *I* with protein *j*. Several proxies of a single group-covariance measure exist. For practical purposes, the average of the lower triangular entries of the correlation matrix was used as a proxy of covariance.

## RESULTS

### Random Forests can detect protein complexes in simulated organelle proteomics data

Proteins in multi-protein complexes have been shown to covary across quantitative proteomics experiments of organelles (8, 9). That is, the absolute or relative quantities of proteins that together form a complex increase or decrease in a coordinate manner. This concerted behaviour forms a potentially detectable ‘signature’ of the complex across sets of proteomics experiments. Other proteins that share the same signature may be functionally related to the complex.

We wondered how strong such a signature would need to be for its detection. The signature is an outcome of the resemblance of each protein’s behavior to each other and how much the group stands out from other groups. We reasoned that the strength of the signature could be modulated in two ways: a) by controlling the fraction of informative experiments (experiment subsets where the members of the complex correlate) and b) by different amounts of noise. Less informative experiments should ‘dilute’ the complex’s signal, whereas stronger noise would lead to fluctuations away from the common behaviour. We therefore constructed artificial proteomics data in which we could independently control these two properties and evaluate their influence on detecting a hypothetical complex.

We generated artificial proteomics tables (Figure 1A) by populating random values into tables of 20 ‘experiment’ columns by 5000 ‘protein’ rows. In those tables, 12 ‘proteins’, which were intended to represent a hypothetical protein complex, were constrained to be identical in a fraction X of columns, while leaving independent random values in the remaining experiments. This action imitated situations in which a complex covaried in only an informative subset of experiments (Figure 1A, middle panel). Next, we jittered all the entries in the table by adding Gaussian noise of strength Y. Figure 1B illustrates the data generated by this approach and exemplifies visually how the number of informative variables and noise contribute to a protein group’s signature behaviour.

We wondered first if the mean of pairwise correlations between proteins of a complex would suffice to reveal membership as levels of noise and informative experiments changed. As one would expect, when the noise was low and the fraction of informative experiments was high, protein correlation was high. However, it dropped rapidly with slightly weaker signatures (Figure 1C).

We then asked if the machine learning algorithm “Random Forests” would recognise stronger or weaker signatures in the behaviour of the hypothetical complex (for an introductory explanation of the algorithm, see methods). Specifically, we asked whether the algorithm Random Forests could distinguish our hypothetical complex from >350 other proteins, composed of >350 rows in the random protein table (Figure 1A, middle panel). In two previous works from our group (1, 9), we used Random Forests because it a) samples combinations of experiments and attempts to draw a ‘boundary’ between a positive and negative class, b) does not make any assumptions about the data, c) can handle missing values, and d) For every ‘protein’, RF outputs a score between 0 and 1 – the RF-score – indicating whether the ‘protein’ behaves as being part of the hypothetical complex (20, 21). Proteins part of the positive and negative classes also obtain an unbiased score regardless of their membership to the training classes (see methods).

Figure 1 D shows that the RF score of the hypothetical complex remained high even with few informative experiments, but fell significantly with higher noise. Therefore, if looking at the RF score alone, even small amounts of noise could lead to not recognising members of the true complex (false negatives), even when they initially had a fairly strong correlation. These results suggest that the RF score is, on its own, not robust to noisy data even when correlation in a complex is high.

We reasoned that a noise-induced decrease in RF scores could be tolerated as long as the scores of members of the hypothetical complex were overall higher than those of the negative class. Yet, levels of noise too high, and too few informative experiments, could lead to false positives. To strike a balance, we searched for a RF score that, if used as a boundary between the two classes, maximized separation quality – i.e. made the fewest class misassignments – between the hypothetical complex and the hypothetical contaminants. This can be assessed by the Matthews Correlation Coefficient (MCC - Exemplified in Figure 2A, lower panels). Figure 1F shows that class separation quality remains for different levels of noise and a small fraction of informative experiments. All measures showed the lowest values for the weakest signatures, where the complex can no longer be distinguished from randomly covarying groups. Altogether, we conclude that RF is able to distinguish significant signatures of a protein group in high noise and few informative experiments, even though the group could be as small as a protein complex. Because of the small training set size, we refer to this instance of Random Forests as nanoRF.

**Figure 2.**
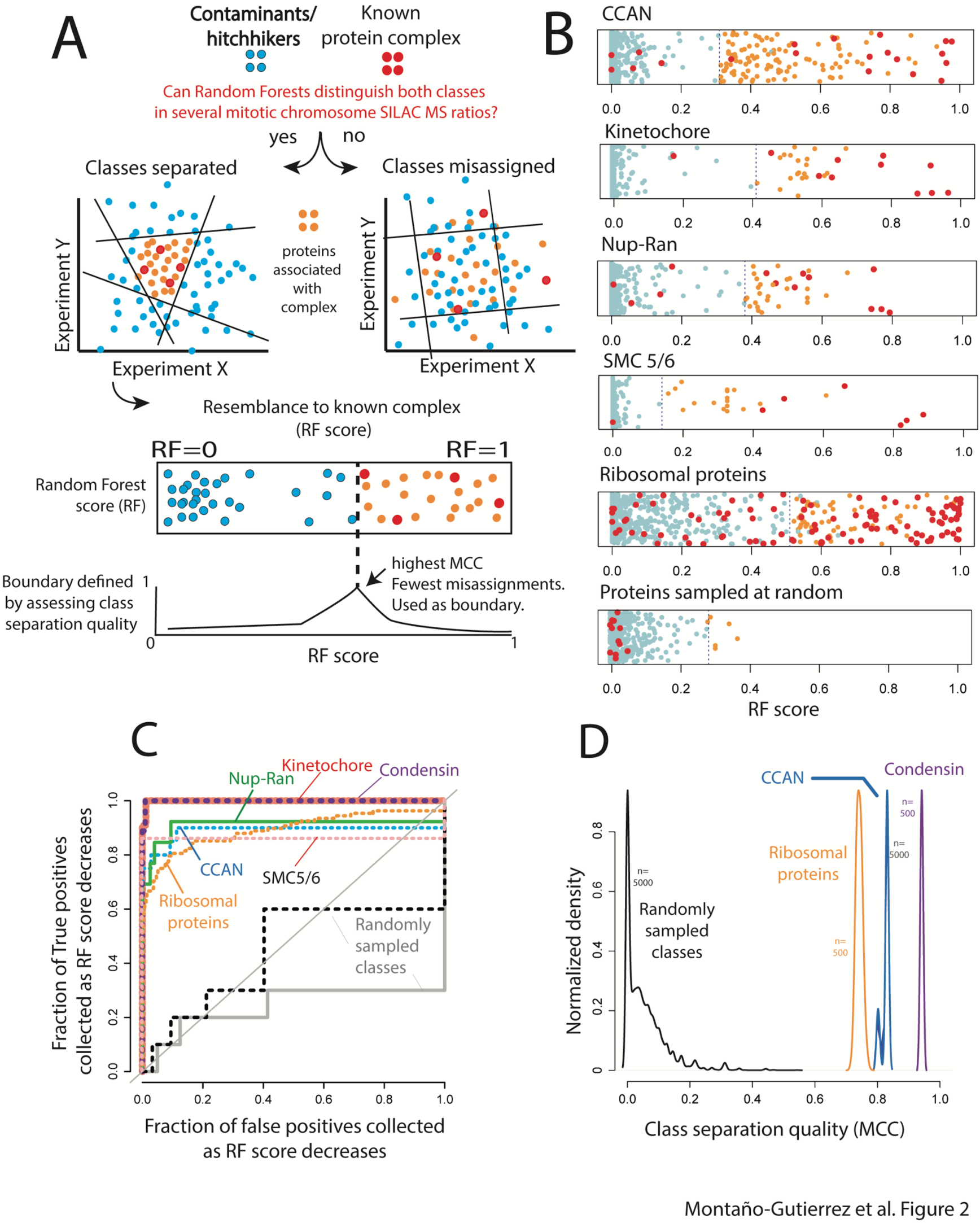
Random Forests can detect small protein complexes in chicken chromosome SILAC proteomics experiments. Entire figure: red-protein complex, blue tones-contaminants/hitchhikers. A. Logic of the procedure to detect complexes with Random Forests. Groups separable in multiple dimensions (only 2 depicted) yield a higher MCC than inseparable groups. B. RF scores of multiple complexes versus the same set of contaminants/hitchhikers, and randomly selected groups from the table. C. Receiver operating characteristic (ROC) performance curves of the RF as a classifier for each protein complex and for two randomly selected protein groups (grey, black). Diagonal shows the random assignment scenario. D. Kernel densities of MCC values for 500 random forest runs of each complex and 5000 runs for randomly assigned groups (black. Sample sizes: 10 for positive class and 425 for the negative class). All distributions were made of height 1 for visualization purposes.

### RF can distinguish protein complexes from contaminants in proteomics experiments of mitotic chromosomes

Our group has both collected and published SILAC proteomics data of mitotic chromosomes isolated from chicken DT40 wild type and knockout cell lines. The proteins targeted for knockouts belong to a range of mitotic chromosome complexes of two groups: Structural Maintenance of Chromosomes (SMC complexes, like condensin SMC2-4., cohesin SMC1-3 (13, 23, 24), SMC5-6 (14, 25)) and the kinetochore (Ska3). We have previously used Random Forests to classify between large groups of ‘true’ chromosomal proteins and potential hitchhikers or contaminants. Given that RF could distinguish small covarying groups in simulated data, we asked whether it could detect known small protein complexes based on real data and if any other proteins shared the signature of the complexes.

The diagram in Figure 2A illustrates our strategy to detect protein complexes in mitotic chromosomes and retrieve proteins that may be functionally linked with them. First, we choose a protein complex (Figure 2, red dots), and a set of curated hitchhikers (Figure 2 blue dots (9), which serve as the negative class (Table S2). Then we use RF to distinguish the complex from the hitchhikers on the basis of our proteomics data. As every protein will get a RF score, we look for a ‘boundary’ score that maximizes class separation quality – i.e. that most members of a complex are above it and the most contaminants below. Proteins above that score covary strongly with members of the complex (Figure 2A and 2B, orange dots). To find the boundary, we use the MCC (Figure 2A, bottom panel) as used in the previous section. A more “traditional” way to evaluate the significance of this result is to consider a hypergeometric test. The higher the enrichment of red marbles on top of the cutoff and the lower the number of blue marbles (higher separation quality), the lower the probability of such draw under an equiprobable hypothesis.

We analysed a number of different complexes with RF (Figure 2B). In particular we performed nanoRF on the Constitutive-Centromere-Associated Network, the KNL-Mis12-Ndc80 (The KMN network), Nucleoporin 107-160/RanGAP, condensin, SMC 5/6 and cohesin and ribosomal proteins. For most complexes, a large number (if not all) of the members have greater RF scores than the contaminants, ensuring high quality boundaries between classes.

To rule out whether the approach could classify any arbitrary protein group to be a complex, we ran RF on 5000 random protein sets from our dataset. The size of those sets (10 random positive class proteins and 400 random negative class proteins) were in the range of the chromosomal protein complexes we investigated, which ranged between 7 and 20. It can be observed that an exemplary random positive class intercalates with the random negative class, resulting in a poor separation quality (Figure 2B, bottom panel). In other words, nanoRF does not support the hypothesis that these arbitrary groups are complexes. This contrasts starkly with the success of separating protein complexes from the negative class (Figure 2B, upper panels). We further evaluated the significance of our results using Receiver-Operating Characteristic (ROC) curves (Figure 2C) and the MCC values themselves (Figure 2D). Starting from the highest RF score, a ROC curve evaluates the fraction of positive class members recovered (true positives) on the vertical axis versus the negative class members recovered (false positives) on the horizontal axis. A ROC curve that climbs vertically is favourable because it means that the RF score is sensitive to the complex. Under these circumstances, the area under the ROC curve (AUC) is larger than 0.5. In contrast, if the RF score contained a poor signal, the positive and negative class would be retrieved randomly. In this case, the ROC curve climbs up the diagonal and has an area of around 0.5. In our analysis, all of the complex-specific RF retrieved roughly 70% of the complexes before any false positives were collected (Figure 2C). All our complexes showed an AUC between 0.9 and 0.999 (Table S2), implying accurate classification. In contrast, ROC curves of the randomly selected groups (examples in Fig. 2C, black and grey lines) remained close to the diagonal.

Finally, we evaluated the distributions of MCC values for real complexes and for randomly sampled protein groups. Quantification of class separation quality by the highest MCC value obtained for the random classes was 0.543 (P≈0.0002, N=5000), whereas the minimum MCC value for the complexes’ separation was 0.71 (P≈0.002, N=500). Altogether, these results support the hypothesis that the RF can distinguish between protein complexes and contaminants in real data. Thus, the performance of real complexes is likely the result of biological relationships, rather than an artefact of machine learning. Strikingly, no particular experiment was aimed at studying the Nup107-160/RanGAP complex or ribosomal proteins. This suggests that this biological information is protein complex covariance as previously observed in other works (8, 9) and suggested by the simulations in the previous section. The full list of proteins associated with each complex can be found in table S2.

### Integration of several complex-specific RF reveals known and novel interdependencies between protein complexes

The covariance of each complex could be its unique signature or could overlap with that of other complexes, possibly implying conditional interdependency among complexes. We decided to test this hypothesis with kinetochore subcomplexes as there is significant contact among them. To this aim, we analysed 2D plots of RF for different complexes (Figure 3).

**Figure 3.**
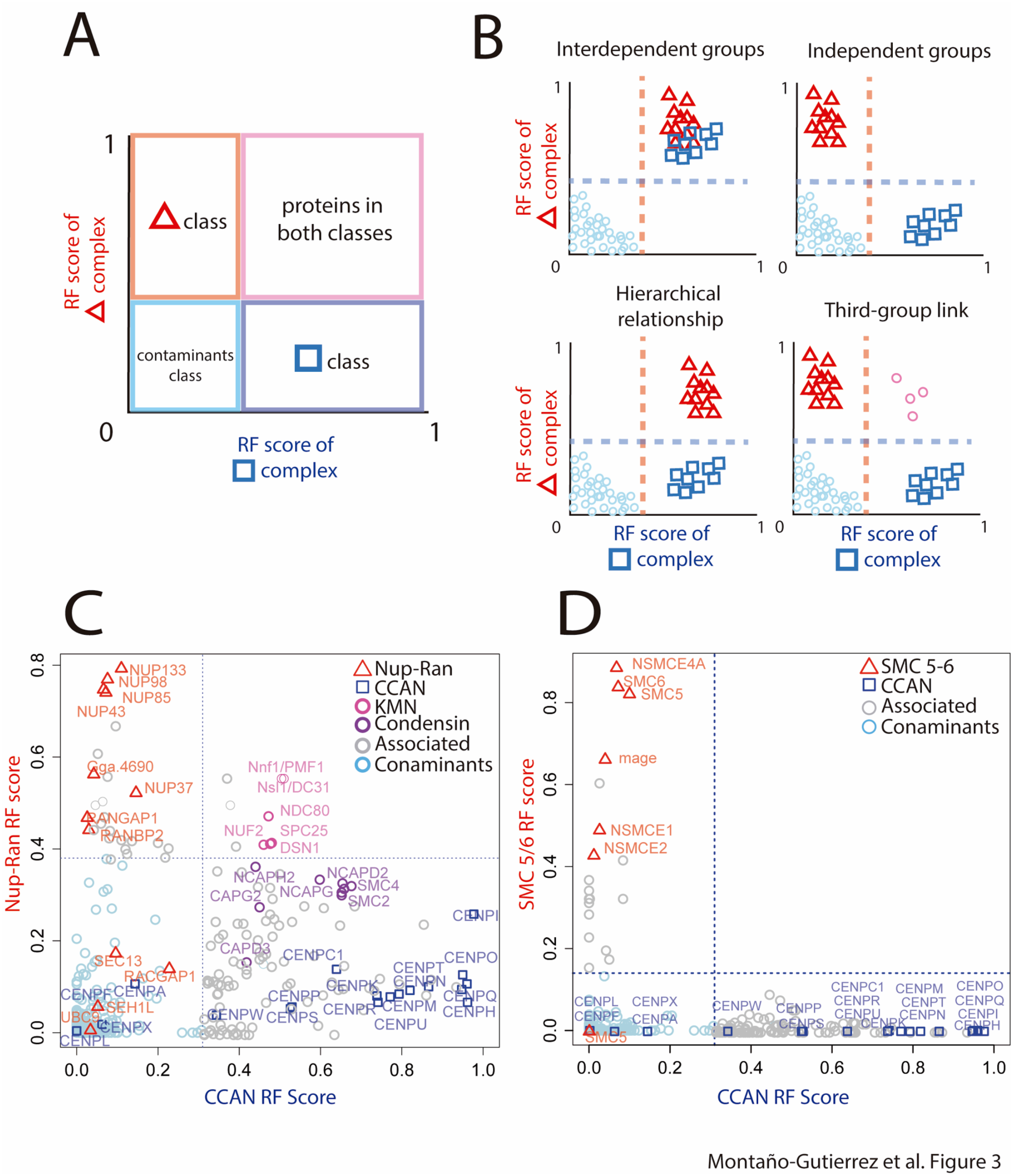
Known and novel interdependencies between complexes revealed by RF. A. Schematic of a 2D diagram to visualize intersections between Random Forests for different complexes. Highest separation quality thresholds are depicted by dotted lines. Proteins above both thresholds (pink quadrant) associate with both complexes whereas those just above one remain independent. B. Possible scenarios of interdependence between complexes inferred from 2D RF plots. C,D. 2D interdependence plot of the Constitutive Centromere-Associated network (CCAN, C and D, squares) versus the Nup107-160/RanGap complex (C, triangles) and the SMC 5/6 complex (D, triangles).

We categorized several possible interdependency scenarios between kinetochore complexes (Figure 3A, B). According to these scenarios, the CCAN and the Nucleoporin 107-160 /RanGAP complex (Figure 3C) appeared independent, i.e. they do not associate with each other. In contrast, the KMN network associated with both. We concluded that perturbations on both CCAN and Nup-107-160 have a hierarchical effect on KMN (i.e. their effects propagate to KMN but not vice versa), implying that the latter is involved in links between inner and outer kinetochore. These observations are consistent with current models of the kinetochore (26, 27). The other proteins associated with the CCAN, Nup-Ran or SMC5-6 complexes can be found in Figure S1.

Even though the CCAN RF prediction was rich in associated proteins – this might be expected from a crowded chromatin environment – the entire condensin complex associated with the CCAN. This dependency may imply a potential relationship between these complexes that merits further study. Finally, Figure 2C shows that the CCAN RF prediction is independent from the SMC 5/6 complex, and no CCAN protein co-fractionated with ribosomal proteins (Figure S2). Together, these results show that, by integrating the outcome of several complex-specific Random Forests, we can reconstruct known dependencies at the kinetochore and identify novel inter-complex dependencies. Notably, none of these relationships were directly addressed a priori by the experiments used.

We suggest that this strategy to infer protein functions and relationships training RF with small protein complexes be named nanoRF. Other sub-complexes and uncharacterized proteins also associated with the complexes shown here. An experimental analysis of putative interactions identified by nanoRF, in the context of SMC complexes, can be found in (11).

## DISCUSSION

A recurrent goal in the post-genomic era has been to make sense of increasing amounts of underexploited data, including noisy and incomplete proteomics output. Our results show that, even with high noise and when few experiments are informative, small groups of covarying proteins–i.e. complexes– can be recognised based on their coordinated behaviour by Random Forests (Figure 1 and 2). In data of this type, statistical measures such as the mean correlation (Figure 1C) or absolute RF score of members in a complex can drop considerably (Figure 1D). We have demonstrated that lower RF scores can be informative as long as the negative and positive class remain separable by their RF score (Figure 1F). By tolerating a decrease of the RF score and maximizing separation quality, we were able to predict highly specific associations with complexes (Figure 2B) and retrieve known inter-complex relationships in our dataset (Figure 3). As no experiment targeted all of the complexes detected, this strategy could potentially identify protein function in any combination of comparable proteomics results.

### Comparison between nanoRF and other methods

Two previous studies from our group, MCCP and ICP, have used Random Forests to attempt to find general trends shared by functional members of chromosomes (9) or interphase chromatin (1) in proteomics data. The evidence presented in the current work suggests that the ‘true chromosome class’ is the integration of the signatures of multiple protein complexes covarying in specific, distinguishable ways. Because of strong, yet conditional complex-specific covariance, adding more than one complex to a training class may restrict the performance of RF. Compared to MCCP and fractionation profiling (11), our prediction would upgrade, for example, from “true chromosomal protein” to “protein dependent on complex A but not complex B”. In a previously unmentioned example, the polybromo-and-BAF-containing (PBAF) complex (ARID2, PBRM1, BRD7, SMARCB1 and SMARCE1) associated specifically with Nup107-160 but not with the CCAN (Figure S1A). In support of this prediction, another bromodomain-containing protein, CREBBP, has been found to interact with Nup98 in Nup107-160 complex and was linked to Nup98 oncogenicity (26).

Methods like Fractionation Profiling (FP) and multivariate proteomic profiling (MVPP) (8) are based on guilt-by-association analyses to similarly detect protein complexes and have cleverly dealt with the intricate nature of proteomics data-i.e. presence of missing values– but the conditional covariance of the complex-i.e. a signal present in only a few experiments– has not been accounted for previously. We have shown that nanoRF finds such covariance, even when there is high noise. Consequently, nanoRF has successfully predicted proteins with previously uncharacterized links to mitosis (11).

### Potential pitfalls and statistical considerations of nanoRF

It is not possible to conclude from computational analysis alone that the relationships predicted by nanoRF are direct physical interactions between the aforementioned protein complexes. Nevertheless, our results come strictly from protein-level dependencies (or indirect effects of these) rather than changing expression levels, so physical associations are likely.

We believe that finding the objectively best separation quality lessens the burden to select an arbitrary significance cutoff for candidates, especially as more uninformative experiments are collected. We have intentionally avoided using a hypergeometric P-Value as a significance measure since a) the exact P-values we obtained for all of our complexes were in the range of 10 ^−11^ to 10^−51^ (Table S2), b) P-Values were strongly influenced by the number of proteins in the complex, c) were undefined for some some of the random group RF results, where none of the two classes were above the MCC threshold (Figure 2B, lowest panel).

Instead of direct P-Value usage, the significance of the predictions by nanoRF is subject to the probability of obtaining a high separation quality by chance for a given dataset. To minimise the risk of type I error, we suggest that the MCC at the classification threshold for a complex remains higher than the highest MCC obtained from randomly assigned protein groups in a data set. In our analysis, the probability of obtaining an MCC as high as that of real complexes by chance showed negligible-our sampled MCC distributions did not overlap (Figure 2D), but it may vary for other datasets. Naturally, a lower MCC may be accepted at the risk of more false positives.

For prediction of associations with a complex, the false discovery rate for each complex should be proportional to the fraction of negative-class proteins that surpass the classification threshold. A small negative class could lead to underestimating false positives as higher noise may increase the RF score of spurious proteins. Therefore, a large negative class may be essential for a realistic False Discovery Rate estimation (28) and a small one could be compensated with a more stringent prediction cutoff for the RF-score.

### Potential applications of nanoRF

In the context of all the massive protein-protein interaction networks being identified, we face a lack of detail in the functionality, hierarchy, specificity and conditionality of these interactions. We have shown that nanoRF could satisfy these unmet needs by providing deep insight about protein complexes.

Experiments are informative if members of a complex covary in them (Figure 1A). Differentiating between informative and non-informative experiments (feature selection) could itself be a powerful tool for protein complex data mining. For example, a specific set of perturbations may break the stoichiometry (and hence the correlation) in a complex. In this direction, our nanoRF pipeline (22) includes a calculation of each experiment’s ‘importance’ for classification, though exploiting such importance may not be straightforward. This estimation employs the Gini importance, which compares classification performance with or without a given experiment. A thorough analysis of importance measures is provided by (29).

We speculate that nanoRF could be performed on the same complex multiple times, each time using a distinct subset of experiments. These subsets could correspond, for example, to different time points or biological conditions, such as drug treatments. Such analysis could potentially inform how the capacity to retrieve a complex changes with the experiments, or whether there is a difference in associated proteins from one condition to the next. Such changes in retrieval may provide insight about conditional binding partners, or the biology of specific conditions, drugs or diseases.

## CONCLUSION

Here we described NanoRF, which uses supervised machine learning to a) detect protein complexes of interest in noisy datasets with few informative experiments, b) predicts proteins that have functional associations with specific complexes and c) evaluates the relationship between complexes according to their behaviour. NanoRF enables hypothesis-driven data analysis from ever-increasing, underexploited quantitative proteomics data. It is generally assumed that machine learning requires large training sets to work. However, we have established that Random Forests can retrieve strikingly small protein complexes, their associated proteins and relationships between complexes from ordinary proteomics data. We anticipate nanoRF to complement experimental co-fractionation approaches such as immunoprecipitation. Importantly, nanoRF does not require proteins to remain physically attached to each other during analysis, which may be difficult for weakly interacting or insoluble protein complexes such as associated in chromatin or membranes.

## Acknowledgments

This work was supported by a Wellcome Trust four-year studentship [grant number 089396] to LFM, a grant from the Uehara Memorial Foundation and the Nakajima Foundation to SO, a Wellcome Trust Senior Research Fellowship [grant number 103139] to JR and a Wellcome Trust Principal Research Fellowship [grant number 107022] to WCE. The Wellcome Trust Centre for Cell Biology is supported by a core grant [numbers 077707 and 092076] and the work was also supported by Wellcome Trust instrument grant 091020.

---

**Figure S1.**
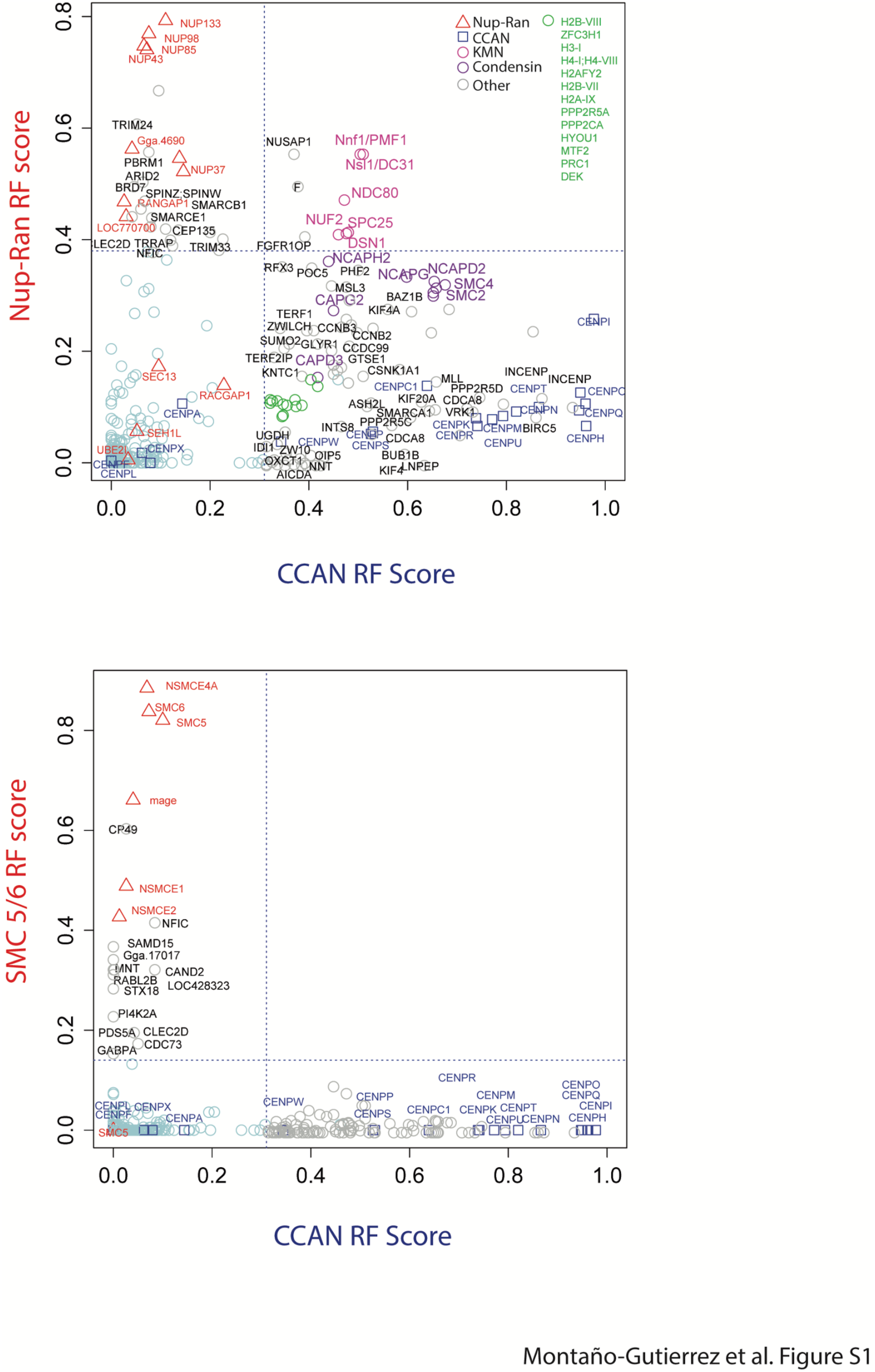
Expanded version of 2D interdependency plots in Fig 3C(A) and 3D(B) shows proteins with functional association to either complex. A. Expanded version of 3C. Green list corresponds to proteins (histones) in green circle cluster. B. Expanded version of 3D. Names were slightly moved to avoid overlap.

**Figure S2.**
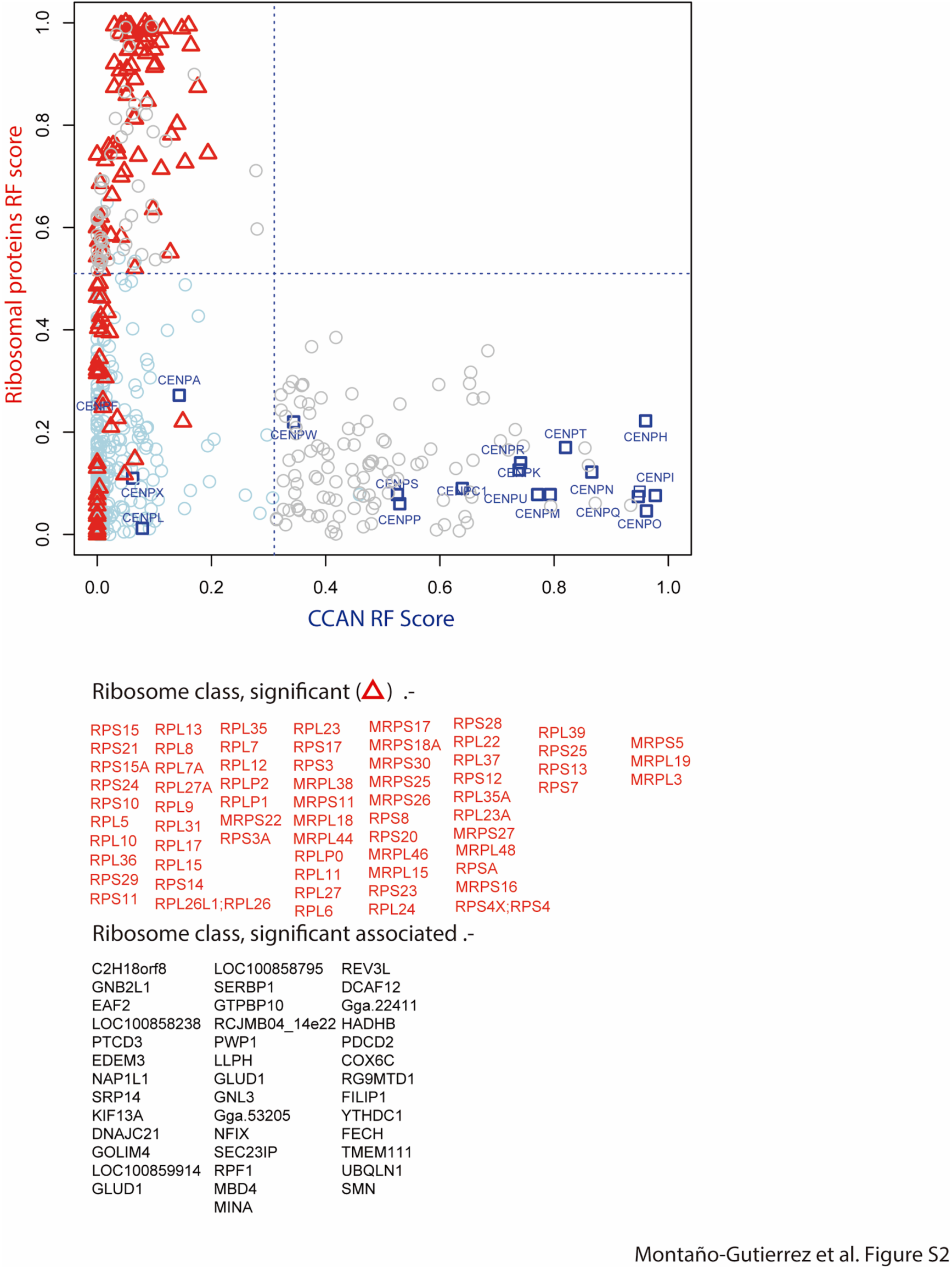
2D interdependency plot between the Constitutive Centomere-Associated Network (CCAN, X axis, squares) and the ribosomal protein group (Y axis, triangles).

**Table S1. nanoRF proteomics results table**. Grey columns: the training factors used for each complex’s nanoRF where T= member of a complex, ‘F’= hitchhiker, and ‘?’= unknown, proteins that are uncalled as any specific class. Orange Columns: the SILAC ratio columns used from the mitotic chromosome proteomics experiments, b) Colourless columns: each RF scores for each complex. Red-coloured entries are proteins that surpass each RF’s cutoff score-i.e. they are significantly associated with the complex. Table S2. Information about protein complexes and associations. Relevant information about the complex composition and statistics (MCC, AUC, RF cutoffs, hypergeometric tests).

